# Track-Control, an automatic video-based real-time closed-loop behavioral control toolbox

**DOI:** 10.1101/2019.12.11.873372

**Authors:** Guang-Wei Zhang, Li Shen, Zhong Li, Huizhong W. Tao, Li I. Zhang

## Abstract

Approaches of optogenetic manipulation of neuronal activity have boosted our understanding of the functional architecture of brain circuits underlying various behaviors. In the meantime, rapid development in computer vision greatly accelerates the automation of behavioral analysis. Real-time and event-triggered interference is often necessary for establishing a tight correlation between neuronal activity and behavioral outcome. However, it is time consuming and easily causes variations when performed manually by experimenters. Here, we describe our Track-Control toolbox, a fully automated system with real-time object detection and low latency closed-loop hardware feedback. We demonstrate that the toolbox can be applied in a broad spectrum of behavioral assays commonly used in the neuroscience field, including open field, plus maze, Morris water maze, real-time place preference, social interaction, and sensory-induced defensive behavior tests. The Track-Control toolbox has proved an efficient and easy-to-use method with excellent flexibility for functional extension. Moreover, the toolbox is free, open source, graphic processing unit (GPU)-independent, and compatible across operating system (OS) platforms. Each lab can easily integrate Track-Control into their existing systems to achieve automation.

## Introduction

Optogenetics empowers us to manipulate neuronal activities in real time during on-going behaviors, which has revolutionized our approaches to functionally dissecting brain circuits. For instance, many labs are using real-time place preference (RTPP) test in which the optogenetic stimulation is controlled based on the spatial location of the animal, to reveal cell-type specific coding of valence in certain brain regions (Carta et al., 2019; Hung et al., 2017; Jimenez et al., 2018; Stamatakis and Stuber, 2012; Xu et al., 2019; Zhang et al., 2018). This behavioral test is easy to apply, and with a collaborative effort, the neuroscience community may soon establish a framework for valence coding in the entire brain.

The application of event-triggered feedback control is however not limited to RTPP. For example, location-based presentation of threating visual stimuli can be used to study innate various types of defensive behaviors (Evans et al., 2018). Stimulation triggered by head orienting towards goals has been used in brain-machine-interface assisted navigation (Richardson et al., 2019). Optogenetic stimulation triggered by social contacts has been applied to understand the rewarding system (Carta et al., 2019). And optogenetic stimulation triggered by food intake has been used to understand feeding circuits (Moreira et al., 2019; Musso et al., 2019). Moreover, online feedback control can provide a therapeutic strategy in disease management. For example, ankle position triggered optogenetic stimulation has been studied to potentially restore motor function in paralysis (Srinivasan et al., 2018). Neuronal activity dependent electrical stimulation has been used to suppress seizures (Grosenick et al., 2015; Srinivasan et al., 2018).

Here we introduce the design and application of Track-Control, a free and opensource toolbox used in our published research (Zhang et al., 2018), which automatically detects spatial coordinates of the animal and sends feedback control to achieve optogenetic stimulation. We have further optimized and upgraded the Track-Control toolbox so that it can be applied in a broad spectrum of standard behavioral tests such as open field, elevated plus maze, Morris water maze, social preference tests, etc. Track-Control can achieve realtime detection of triggering events (e.g. specific locations of the animal, static or dynamic) and output statistical results immediately after the behavioral test is complete.

The Track-Control toolbox is video-based and written in Python programming language (compatible with Python 2 and Python 3). It works on Windows, Max OS and Linux system. Due to low computational loads, Track-Control can run on a single core CPU computer for real-time object detection and feedback control. Considering that most neuroscience labs have laser/LED components for optogenetic stimulation or customized systems for sensory stimulation, the Track-Control toolbox can be easily integrated into existing laboratory setups to achieve automation. The source code and detailed tutorials can be accessed from https://github.com/GuangWei-Zhang/TraCon-Toolbox/.

## Results

The Track-Control toolbox can achieve real-time object detection and send low latency feedback control to another hardware through a USB port (**Fig. 1**). The toolbox is based on the OpenCV library (https://opencv.org), consisting of an object detection part and a feedback commanding part. From our own experience, one-for-all software may not be so convenient for behavioral labs, especially considering that one computer is often assigned to a specific type of behavioral test. Thus, we decided to distribute an array of python modules, from which only a specific one is used for a particular behavioral test. The control settings and analysis methods have been optimized for each individual test, and the module is ready to use without further modifications.

**Figure 1.**
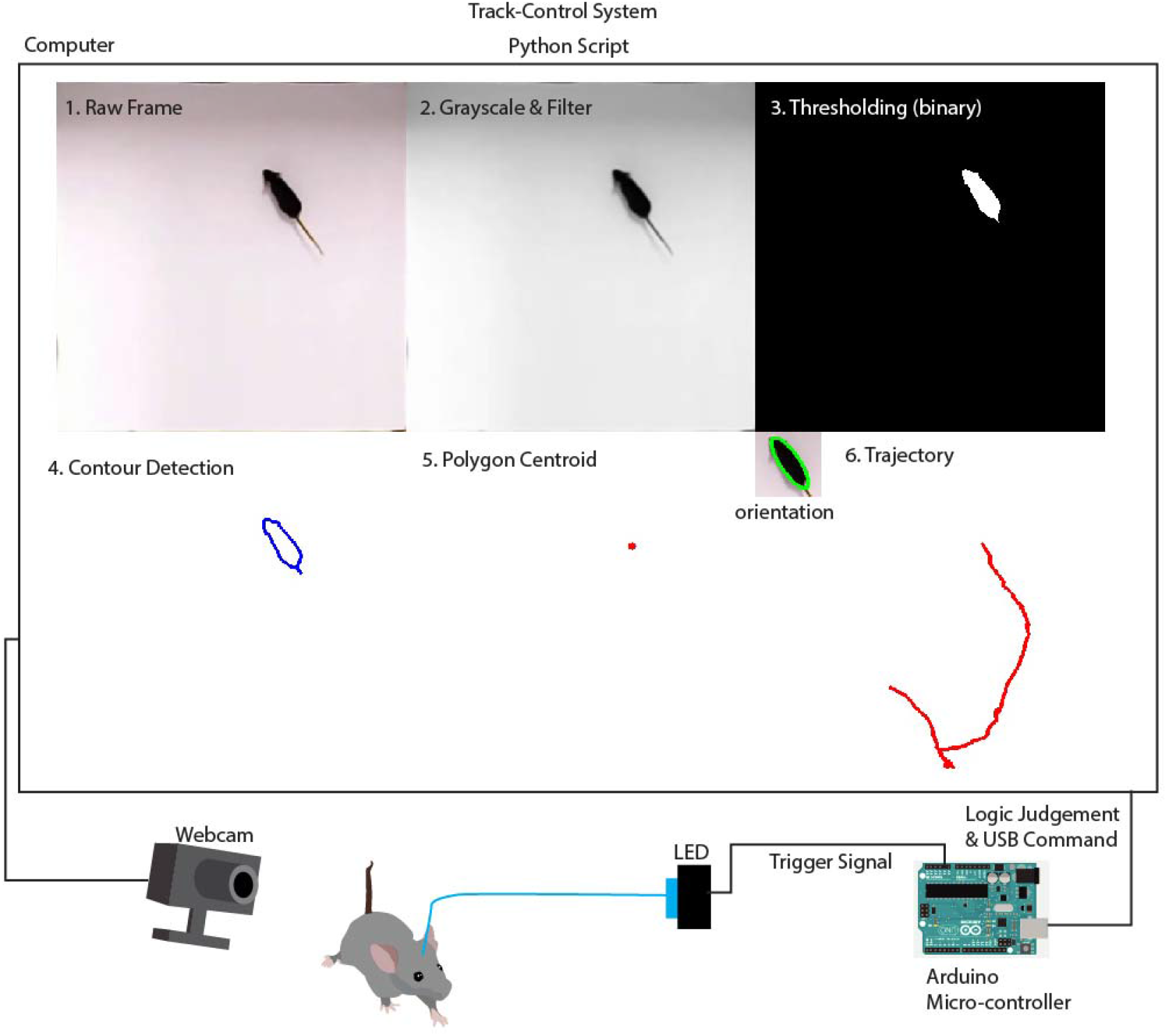
Structure of the Track-Control system. Schematic illustration of the Track-Control system. Images from the webcam are streamed into the computer. Each frame is first converted to a grayscale image. After gaussian filtering and binarization, animal contour in form of a polygon is detected and the centroid of the polygon is determined. Logic judgment is then performed to determine whether a command signal will be sent via the USB port. The signal is used to trigger hardware output (e.g. LED light) via an Arduino micro-controller.

### Object detection

Track-Control initiates and controls the video acquisition, and processes each frame streamed from a webcam (**Fig. 1**). In the meantime, the video file is stored to the hard drive of the computer while performing the experiment and the default output file format is .mp4. Most brand webcams can satisfy the frame rate requirement for the majority of behavioral tasks (up to 30Hz). Here, we use the default frame rate of the webcam used.

The pipeline for each frame is first to convert the RGB image to greyscale, by averaging the value of the three-color channels. Then a 2D Gaussian filter is applied to suppress high frequency noise in the image:

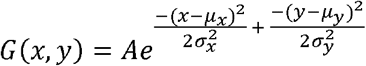

with σ, μ = 5 in the kernel. Binary thresholding is next applied to generate binary images and separate the foreground from the background:

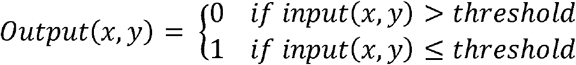

Next, we extract the object contours based on a border-following algorithm (Suzuki and be, 1985). The centroid of the extracted polygonal contour is used to define the position of the animal, and the x and y coordinates in each frame are stored in a .csv file for further analysis (**Supplementary Movie 1**).

For enhancing the contrast between the object and the background, we suggest using a light color background (e.g. white) for pigmented animals (e.g. C57BL/6 mice), and dark color background for albino animals (e.g. FRT). The detection accuracy of Track-Control is comparable to existing offline detection methods (e.g. DeepLabCut), but Track-Control can run much faster on a single core CPU computer. The software can perform both online analysis and offline analysis of pre-recorded video files, with compatibility depending on the installed video encoder. Here, .avi, .mp4 and .mov file formats have all been tested and Track-Control can handle them well.

### Trajectory overlay and data analysis

Upon detection of animal location in each frame, the trajectory of movements is plotted and overlaid on the video, facilitating visualization of the locomotion pattern (**Supplementary Movie 1**). Besides data visualization, Track-Control can provide quantitative measurements of the animal movement. Here, we demonstrate that the tracking module of Track-Control can be applied in an open field test (**Fig. 2A**), which has long been used for testing anxiety. Based on the conflict theory, animals with higher levels of anxiety would spend more time in the peripheral (including corners) than the central region of the test arena where there is a higher risk of exposure to potential predators (Calhoon and Tye, 2015; Tovote et al., 2015; Tye et al., 2011). Thus, the percentage time spent in the central region is quantified. Boolean value based on the spatial location of the animal for each frame is returned sequentially:

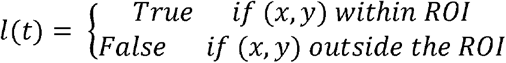

**Figure 2.**
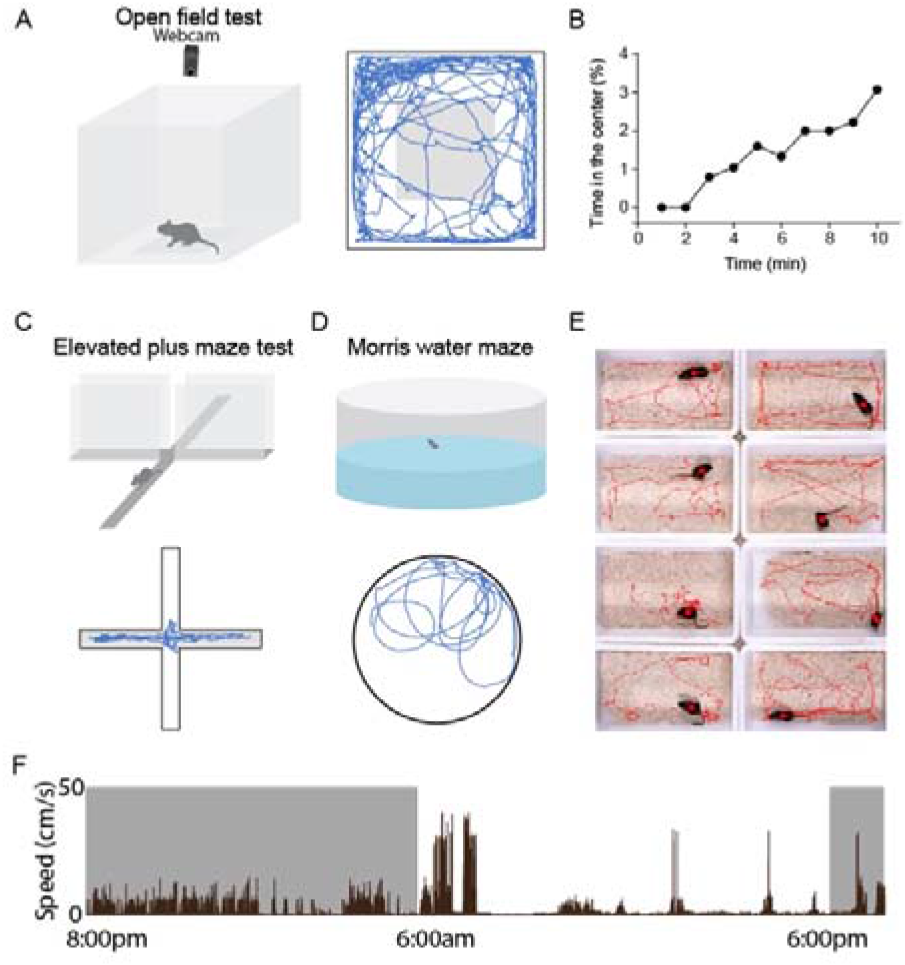
Animal detection and trajectory plotting in behavioral assays. (**A**) Left panel, schematic illustration of an open field test. Right panel, locomotion trajectory for an example animal, with the gray box representing the center region. (**B**) Percentage of time animal spent in the center region, calculated over each time bin, for the same animal shown in (**A**). (**C**) Top panel, illustration of an elevated plus maze (EPM) test. Bottom panel, trajectory of an example animal in the test, with gray rectangles representing closed arms. (**D**) Top panel, illustration of a Morris water maze test. Bottom panel, trajectory of an example animal. (**E**) Simultaneous recording of 8 test boxes using one webcam. Trajectories of 8 animals are simultaneously plotted. (**F**) Plot of locomotion speed for an example animal during 24-h surveillance. Grey box represents dark cycle.

And the percentage of time in a designated region of interest (ROI, such as center of the arena) is simultaneously updated:

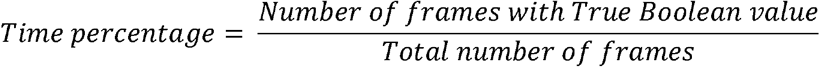

Here, we compared the anxiety level between an autism animal model (16p11.2df, which harbors a chromosome microdeletion) and normal animals. The autism model mouse shows elevated anxiety compared to normal animals (**Fig. 2B**). The tracking module can also be easily applied to elevated plus maze (EPM) test (**Fig. 2C**) to calculate the percentage time spent in the open arms of the maze, which is also an indicator of anxiety level. In Morris water maze test (**Fig. 2D**), we can calculate the travel distance and time spent in each quadrant. Moreover, Track-Control can be applied simultaneously to multiple spatially nonoverlapping test areas, which greatly expands throughput (**Fig. 2E**). The serial coordinates of the animal and Boolean values for all frames are stored in a .csv file that can be used for deeper data mining.

### Long-Term Tracking

As discussed in a previous study (de Chaumont et al., 2019), long-term recording can be a problem for most available tracking programs, especially those requiring manual interference. Here, we demonstrate a 24-h recording and tracking with Track-Control without needs for any manual interference (**Fig. 2F**). In principle, the recording length is only limited by the volume of hard drive storage. Such chronic tracking can potentially be used to investigate the circadian rhythm (**Fig. 2F**), as an alternative to running-wheel assays (Verwey et al., 2013).

### Online Feedback Control

To achieve a closed-loop feedback control, the Boolean value will feed to an external device through the USB port. In Track-Control, users may define a trigger event, for example, when animal enters a designated location (i.e. Boolean value = True). The control signal for feedback stimulation is relayed by an Arduino Microcontroller (UNO board) to a stimulation device such as an LED driver.

We measured the latency of the system using a high frame rate camera (240fps). The latency is defined as the time lag between the moment animal enters the designated location and the onset of LED light. We found that the time lag was 1 frame (i.e. < 4 ms) (**Supplementary Fig. 1**). Therefore, the feedback stimulation can almost be instantaneously triggered.

### Application in looming-induced defense behavior

Looming stimulus such as an expanding dark disc which mimics the shade of an approaching predator can elicit defensive behaviors such as freezing and flight in rodents (Evans et al., 2018; Yilmaz and Meister, 2013). The visual stimulation was manually triggered by experimenters in previous studies (Yilmaz and Meister, 2013). Here, we demonstrate that Track-Control can achieve automation of the visual stimulation with a precise control of visual angle. The test was performed in a rectangular acrylic box, with an LCD monitor placed on top of one side of the box (**Fig. 3A**). The looming stimulation was coded using Psychtoolbox-3 (http://psychtoolbox.org). Before starting the test, users can define the ROI to trigger looming stimulation. By controlling the relative position of the animal to the LCD monitor, we can standardize the visual angle of stimulus at the onset of looming stimulation. The result shows that looming stimulation can robustly elicit an initial freezing (speed decreases to zero) and then a flight (speed rebound) response (**Fig. 3B**).

**Figure 3.**
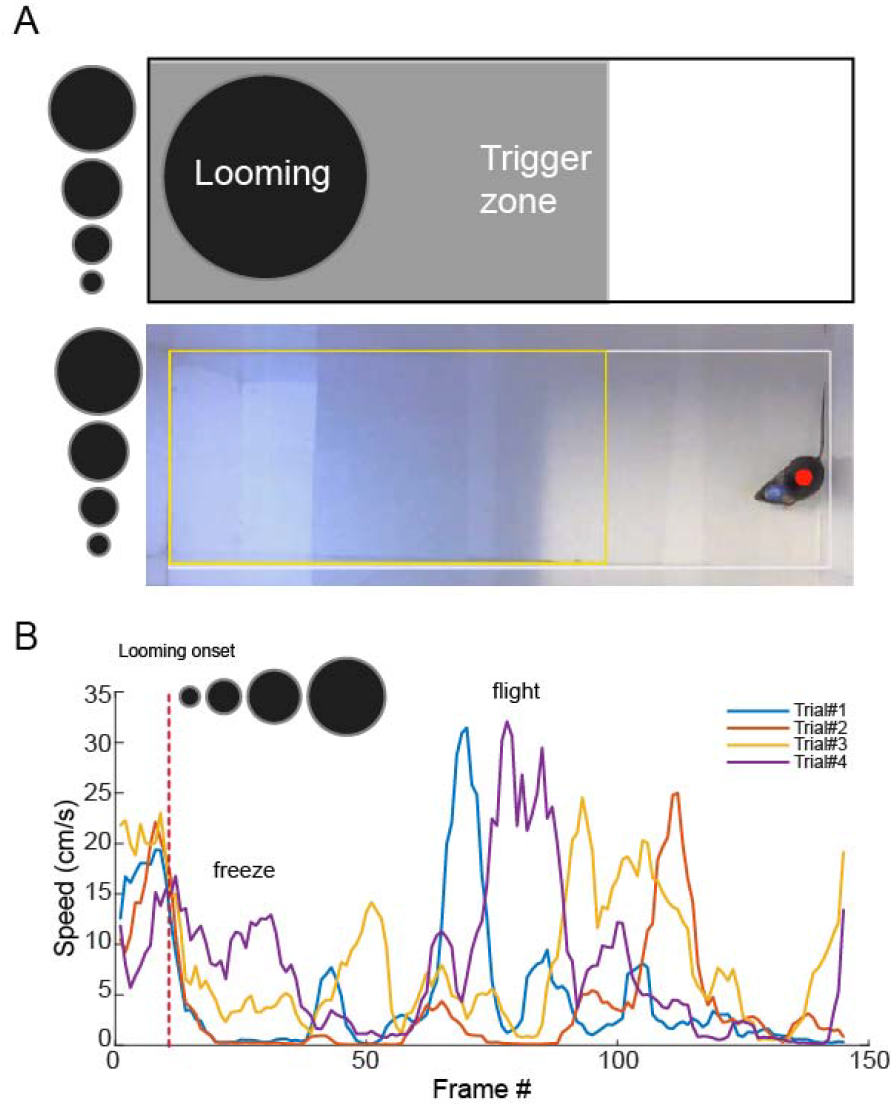
Application of Track-Control in visual looming induced defensive behavior. (**A**) Upper panel, illustration of behavioral setup. Lower panel, a representative image of the behavioral test. White rectangle outlines boundaries of the test field. Yellow rectangle outlines user-defined trigger zone. Red dot depicts the centroid of the animal. When Track-Control detects animal entering the trigger zone, the system triggers the looming stimulation (expanding dark disk from the upper visual field). (**B**) Plot of the speed (aligned by the onset of looming stimulation) for the first 4 trials for an example animal. The animal exhibited first freezing (speed decreased to zero) and then flight (a sudden increase in speed) responses following the onset of looming stimulation.

### Application in real-time place preference test

Besides triggering sensory stimulation, Track-Control can be used to standardize other behavioral tests. Here, we used a real-time place preference (RTPP) test to demonstrate the real-time feedback control using the Track-Control toolbox. Dopaminergic neurons in the ventral tegmental area (VTA) has long been known for its involvement in motivated behaviors (Beier et al., 2015; Matsumoto and Hikosaka, 2009; Roeper, 2013; Zweifel et al., 2011). We tested our Track-Control system in recruiting dopaminergic neurons in the RTPP test. To stimulate dopaminergic neurons, we injected bilaterally adeno-associated virus encoding Cre-dependent channelrhodopsin2 (AAV-DIO-ChR2) into VTA of DAT-Cre transgenic mice (**Fig. 4A**). Two weeks following the virus injection, we implanted optic cannulas into the VTA bilaterally. After one-week recovery, the animal was habituated to the optic fiber connection for 3 days.

**Figure 4.**
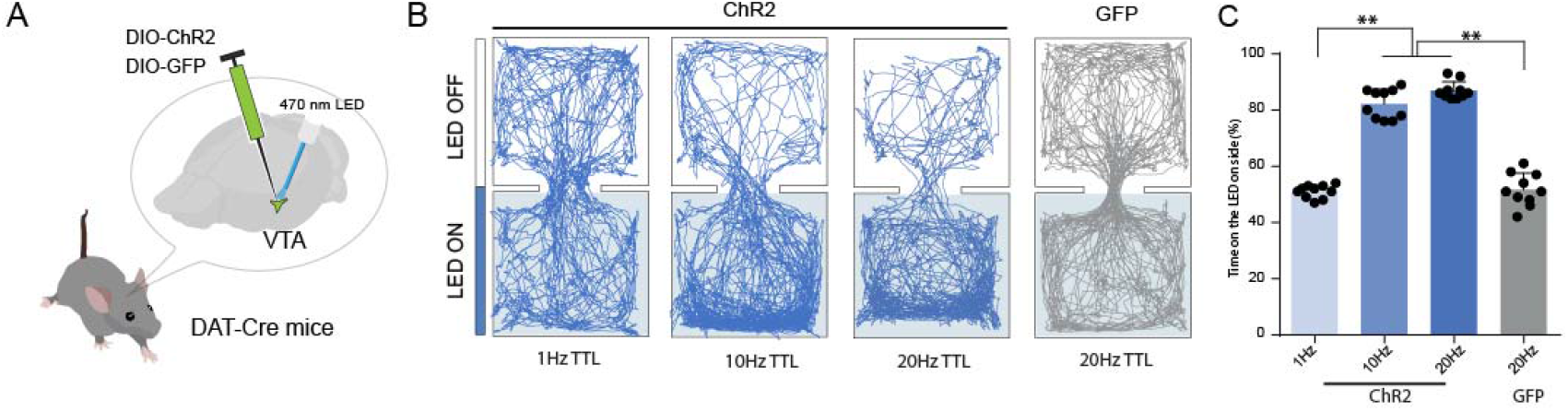
Track-Control application in a real-time place preference test. (**A**) Schematic illustration of viral injection into VTA and optic fiber implantation. (**B**) Locomotion trajectory for a ChR2-expressing animal (blue) under different stimulation frequencies and a GFP control animal (grey). (**C**) Quantification of percentage of time spent in the stimulation (LED ON) chamber. **p < 0.01, One-way ANOVA test. N = 10 ChR2-expressing and 10 GFP control animals.

The RTPP test was performed in an acrylic box divided into two chambers. When the animal entered the chamber designated for stimulation, the Track-Control system sent a command signal to trigger pulsed LED stimulation (at 1Hz, 10Hz or 20Hz stimulation frequencies) until the animal moved out of the chamber (Zhang et al., 2018). Thus, the trigger event was determined by the animal’s coordinate within a defined space, which can be easily adapted to other spatially controlled behavioral tests. We found that testing groups (10 and 20Hz stimulation frequencies, expressing ChR2) spent significantly more time in the stimulation chamber compared to the control group (expressing GFP) or 1Hz stimulation frequency group (ChR2, n = 10 for each group, **P< 0.01, One-way ANOVA, post-hoc test, **Fig. 4B-C**). This result is consistent with the role of dopaminergic neurons in appetitive motivation (Beier et al., 2015; Matsumoto and Hikosaka, 2009; Roeper, 2013; Zweifel et al., 2011).

The stimulation parameter can be defined in the microchip programmed using the Arduino IDE (https://www.arduino.cc/en/Main/Software). When receiving the USB command, a TTL train (e.g. 20Hz, 5ms on, 45ms off) is delivered to a LED driver to achieve optical stimulation.

### Application in social cue-dependent associative learning

In RTPP test, the ROI is pre-defined and static. Next, we demonstrate that the ROI can be dynamic when applied in studying a social interaction behavior. In this behavioral test, two animals are placed in a long corridor but are separated by a transparent acrylic wall with holes in it so that visual and odor cues are not blocked. We defined the ROI as the spatial location where the two animals were close to each other. We expressed ChR2 in VTA neurons (**Fig. 5A**) in one of the two animals and this animal would receive LED stimulation when the two animals were in a close distance (**Fig. 5B**, **Supplementary Movie 2**). We found that over time the average distance between the two animals decreased (**Fig. 5C**). This result suggests that VTA neuron activity promotes social interaction. Thus, we demonstrate that Track-Control can be used when dynamic stimulation contexts are required.

**Figure 5.**
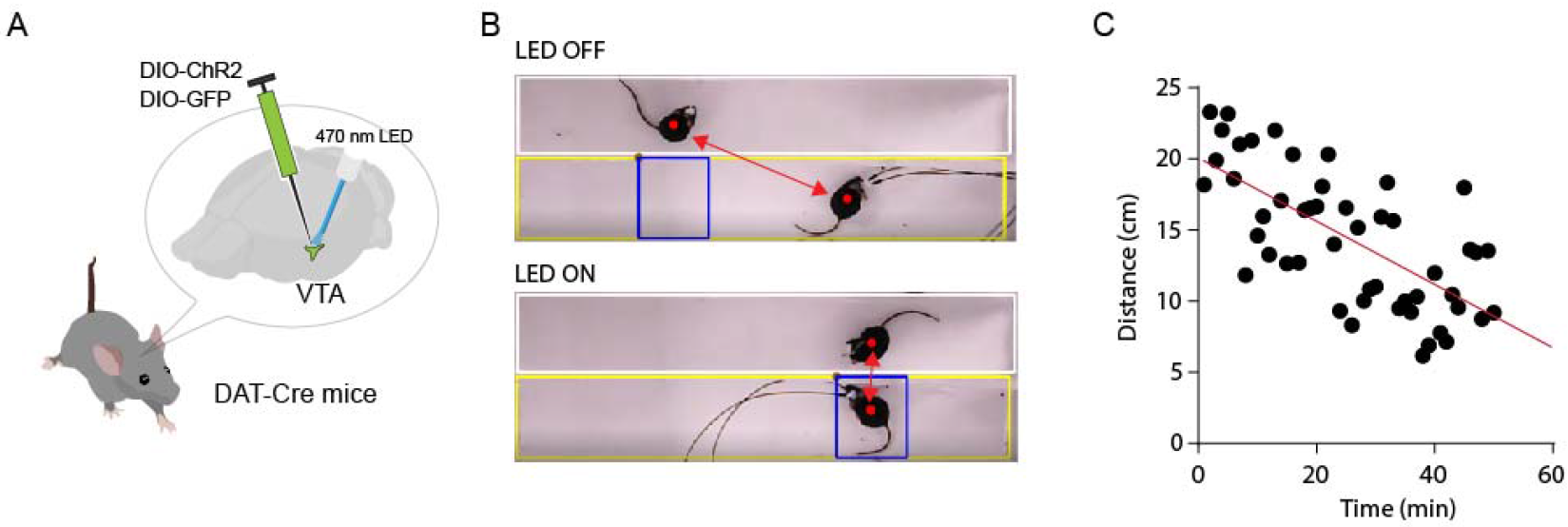
Track-Control application in a social interaction association test. (**A**) Schematic illustration of viral injection and optic fiber implantation. (**B**) Representative images during the behavioral test. White rectangle outlines the corridor of the target animal, and yellow rectangle outlines that of the test animal. Blue rectangle depicts the trigger zone. Red arrows mark the distance between the two animals. (**C**) Plot of average distance between two animals within a 1-min time bin over experimental time. Red line is the linear regression fitting line.

## Discussion

Our goal is to develop an open source toolbox that can detect the position of the animal in real-time and send low-latency feedback control signals. We also intend to control cost so that each lab can build multiple setups to expand the throughput. After testing multiple methods, we found that using contrast and binarization would be a fast, reliable and low computational cost method.

With the rapid development of computer vision and machine learning, object detection has welcomed its great progress and enabled us to automatically analyze videos, increasing the objectivity. For example, DeepLabCut has successfully achieved markerless feature extraction from pre-recorded videos (Mathis et al., 2018; Nath et al., 2019), and another group has demonstrated that the method has potential of being applied in real-time feedback control (Forys et al., 2018). DeepLabCut was built upon ResNet and not focused on the real-time detection. However, fast methods such as YOLO (Redmon and Farhadi, 2018) are available now. In the future, we will implement light-weight and fast network models to facilitate detection and classification of more specific behavioral patterns, e.g. grooming, sniffing and rearing, and to gain a better control in complex behaviors.

Most commercial software packages for object-detection such as ANY-maze (Stoelting Co; IL, USA), TopScan (Clever Sys Inc.; VA, USA) and Opto-Varimex (Columbus Instruments; OH, USA) but except Ethovision® XT (Noldus; Wageningen, The Netherlands) have yet to implement video-based closed-loop hardware control. Even so, the commercial software is still expensive and not flexible enough for functional extension. The neuroscience community has made lots of effort in developing open source software (Gleeson et al., 2017). For example, Open Ephys system has made multichannel electrophysiological recordings flexible and affordable for many labs (http://www.open-ephys.org), which is beneficial for the entire field. Previous animal tracking software programs either require manual interference and are non-open-source (Tort et al., 2006), or are not written using popular programming languages such as Perl (Samson et al., 2015). Some video-based tracking with feedback control methods are not open source yet (Richardson et al., 2019). In comparison, Track-Control is a free, open source and flexible toolbox that provides a strategy for other labs to quickly implement in automatic video-based behavioral experiments.

## Materials and methods

(detailed installation instruction and video tutorials can be accessed at https://github.com/GuangWei-Zhang/TraCon-Toolbox/).

### Animals used in this study

All animal usage and experiment are in accordance with the Animal Care Committee of the University of Southern California. Mice were housed in the vivarium with normal light-dark cycle (light on at 6:00 am and off at 6:00 pm). 8-10 weeks old male C57BL/6 (JAX number: 000664), autism animal model (16p11.2df, JAX number: 013128), DAT-Cre (JAX number:006660) were used in this study.

### Viral tools

AAV1-EF1a-DIO-hChR2-EYFP (Addgene: 35503) was used to modulate the neuronal activity in corresponding regions. AAV1-EF1a-DIO-EYFP (Addgene:27056) was used in control experiments.

### Animal preparation and surgical procedures

We used the following coordinates for injection: VTA: bregma −3.0 to −3.2 mm, lateral 1.25mm at 10 degree-angle, ventral 4.35mm (Hung et al., 2017). Mice were anesthetized with 5% isoflurane and maintained using1.5-2% isoflurane. A small cut was made on the skin covering the craniotomy position and the muscles were removed. One ~0.25-mm^2^ craniotomy window was made for each region. AAVs encoding ChR2 or GFP were used depending on the purpose of experiments and strain of mice. A beveled glass micropipette (pulled using MODEL P-97, Sutter Instrument Co., tip diameter: 15-20□μm) was used to deliver the virus. Virus was delivered by pressure injection and the glass micropipette was attached to a micro syringe pump (World Precision Instruments). For pressure injection, 50 nl of the viral solution was injected at a rate of 15 □nl min. After the injection, the pipette was allowed to rest for 5□min before withdrawal. The scalp was then sutured. Following the surgery, 0.1 □mg/kg buprenorphine was injected subcutaneously before returning the animals to their home cages. Mice were allowed to recover for two weeks. Then, animals were anesthetized with isoflurane and an optic cannula (200 □μm core, NA 0.22, Thorlabs) was stereotaxically implanted into the target region, fixed with dental cement. The mice were allowed to recover for at least one week before behavioral tests. After each experiment, the brain was extracted, sectioned and imaged under a confocal microscope (FluoView FV1000, Olympus) to confirm locations of viral expression and the implantation site. weeks before cannula implantation.

### *In vivo* optogenetic stimulation

During the 3 days before behavioral tests, animals were attached to optical fibers without LED stimulation for habituation. On the test day, the optic fiber (200 □μm core, NA 0.22, Thorlabs) was connected to a blue LED source (480 nm, 0-30 Hz pulses, 5-ms pulse duration, Thorlabs). The LED power measured at the tip of the fiber (connected with optic cannula) was around 3-5mW.

### Behavioral assays

All behavioral tests were performed in the dark cycle of the animal (normally start at 8:00 pm till 2:00 am).

#### Open field test

A white behavior test box (60cm x 60cm x 30cm, length x width x height) was virtually divided into a center region (center, 30 x 30 cm) and a periphery field. For each test, the mouse was placed in the periphery and the locomotion of the animal was recorded by a video camera for 20 min to measure the time spent in the center or peripheral area.

#### Elevated plus maze test

A cross maze with two closed and two open arms was elevated 30cm above the ground. The mouse was placed in the center of the cross maze and the locomotion of the animal was recorded by a video camera for 5 min.

#### Real-time place preference test

A clear acrylic behavior box (40cm x 20cm x 20cm, divided into two chambers, put in a larger white foam box) with normal bedding materials was used. For each trial, the mouse was initially placed in the non-stimulation chamber, and LED (480 nm, 10 Hz, 5-ms pulse duration) stimulation was constantly delivered once the animal entered the stimulation chamber and was stopped when the animal exited. The total duration of each test session was 20 min. Animals were returned to their home cage after each test session. The stimulation chamber was randomly assigned to each animal and balanced for the whole group.

#### Morris water maze test

A circular pool (diameter:100cm) was used, with an invisible platform placed in one quadrant. Water was dyed white using an odorless pigment. Animal was placed in one quadrant. The test was stopped when the animal found the hidden platform.

#### 24-h chronic locomotion test

A sound-attenuation chamber was used, with controlled light illumination (on: 6am; off: 6pm). A continuous infrared illumination was used for observing animal behavior.

#### Social association test

A clear acrylic chamber (40×20×20cm) was equally separated into two corridors (40×10×20cm). The separating wall was transparent and with evenly distributed holes (3mm in diameter) in it so that the two animals placed in the two corridors could see and smell each other. The trigger zone was defined by the location of the object animal and would change its position accordingly. The test animal could freely explore the corridor but only received LED stimulation when located in the trigger zone defined by the object animal. The average distance between these two animals was calculated over 1-min time bins.

### Real-time animal detection and closed loop optogenetic control

Custom mouse detection software was used for online real-time animal detection (written by Guang-Wei Zhang, in Python 3.4, http://www.python.org, with OpenCV library, https://opencv.org) (Zhang et al., 2018). The behavior of the animal was monitored using an infrared camera at 24fps. Then each frame was gaussian blurred and then binarized. The gravity center for the detected contour was used to determine the location of each animal. In the two-chamber place preference test, the stimulation chamber was randomly assigned (balanced within the group) to each animal. Once the mouse entered the stimulation chamber, computer-controlled Arduino microcontroller (http://www.arduino.cc) would generate TTL signals to drive the LED light source (ThorLabs Inc.). The behavior test was run automatically without experimenter’s interference and the result was calculated right after each experiment.

### Looming visual stimulation

The looming stimuli is generated using Psychtoolbox-3 (http://psychtoolbox.org). The looming stimulation was constantly on as long as the animal was detected to be in the trigger zone. Position of the monitor was adjusted to make sure that the visual angle was 75° (from the looming center) when the animal just crossed the boundary of the trigger zone. Diameter of the dark disc changed from 2° to 40° within 250 ms, with a 250 ms interval between sequential presentations of discs.

### Statistics

Pilot experiments were conducted to determine the sample size. One-way ANOVA and post hoc multiple comparisons were used to test significance between samples.

### Data and software availability

The video data were available upon request to the corresponding author. Software and test video files are available on https://github.com/GuangWei-Zhang/TraCon-Toolbox.

### Instructions for software installation

Detailed wiki and step-by-step video tutorials are available on https://github.com/GuangWei-Zhang/TraCon-Toolbox. In brief, first download Anaconda python 3.7 version and set the path as the system default path(https://www.anaconda.com/distribution/). Then in the anaconda prompt command, install the OpenCV (https://opencv.org/) and pyserial module. Arduino IDE can be downloaded from https://www.arduino.cc/en/main/software.

### Arduino coding

Arduino IDE was used to program the UNO board and the script used here can be found in the following GitHub repository (https://github.com/GuangWei-Zhang/TraCon-Toolbox).

## Acknowledgements

The project was supported by grants from the US National Institute of Health (R01DC008983 and RF1MH114112 to L.I. Zhang; R01EY019049 to H.W. Tao).

## Additional information

### Author contributions

G.-W. Zhang designed the Track-Control system and performed all the experiment and analysis and wrote the first draft of the manuscript. L. Shen developed a graphic user interface for an online version of Track-Control. Z. Li designed visual looming stimulation. H.W. Tao and L.I. Zhang supervised the project and edited the manuscript.

