## Supplementary Materials for "Track-Control, an automatic video-based real-time closed-loop behavioral control toolbox"

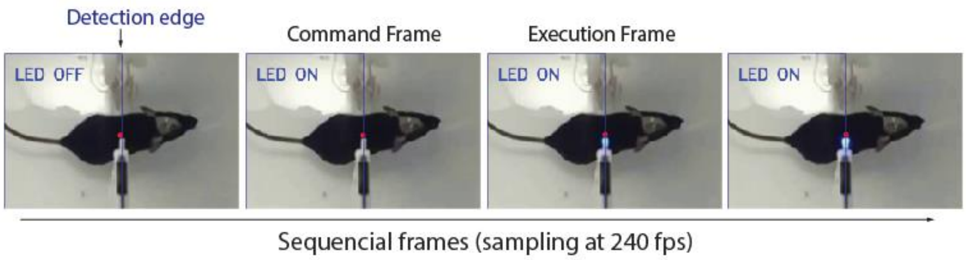


**Supplementary Figure 1.** Measuring latency of the trigger signal.
Four sequential images show the time lag between software judgement and hardware trigger. Upper left corner (LED OFF/ON) shows the software judgement if the detected centroid crosses the defined boundary. Blue line depicted the boundary of user-defined trigger zone. The tip of optic fiber was placed on the side for better visualization of the LED illumination. Using a high-speed camera (240fps), we find the time lag is one frame, which is 4ms.
